# Distribution and localization of phosphatidylinositol 5-phosphate, 4-kinase alpha and beta in the brain

**DOI:** 10.1101/2020.01.29.925685

**Authors:** Evan K. Noch, Isaiah Yim, Teresa A. Milner, Lewis C. Cantley

**Affiliations:** Sandra and Edward Meyer Cancer Center, Weill Cornell Medicine, 413 69^th^ St, New York, NY 10021 U.S.A; Department of Neurology, Weill Cornell Medicine 1300 York Ave, New York, NY 10021 U.S.A.; Feil Family Brain and Mind Research Institute, Weill Cornell Medicine, 407 East 61^st^ Street, New York, NY 10065 U.S.A.; Harold and Milliken Hatch Laboratory of Neuroendocrinology, The Rockefeller University, 1230 York Ave, New York, NY 10065 U.S.A.

**Keywords:** Phosphoinositide, phosphatidylinositol-5-phosphate 4-kinase, PIP4K, brain, neuron, oligodendrocyte, RRID:AB_1127270, RRID:AB_353929, RRID:AB_2164572, RRID:AB_2223210, RRID:AB_10622025, RRID:AB_561049, RRID:AB_10711040, RRID:AB_2269374, RRID:AB_2300649, RRID:AB_1904103, RRID:AB_2096811

## Abstract

Phosphatidylinositol-4,5-bisphosphate (PI-4,5-P_2_) is critical for synaptic vesicle docking and fusion and generation of the second messengers, diacylglycerol and inositol-1,4,5-trisphosphate. PI-4,5-P_2_ can be generated by two families of kinases: type 1 phosphatidylinositol-4-phosphate 5-kinases, encoded by PIP5K1A, PIP5K1B and PIP5K1C, and type 2 phosphatidylinositol-5-phosphate 4-kinases, encoded by PIP4K2A, PIP4K2B, and PIP4K2C. While the roles of the type 1 enzymes in brain function have been extensively studied, the roles of the type 2 enzymes are poorly understood. Using selective antibodies validated by genetic deletion of pip4k2a or pip4k2b in mouse brain, we characterized the location of the enzymes, PI5P4Kα and PI5P4Kß, encoded by these genes. In mice, we demonstrate that PI5P4Kα is expressed in adulthood, whereas PI5P4Kß is expressed early in development. PI5P4Kα localizes to white matter tracts, especially the corpus callosum, and at a low level in neurons, while PI5P4Kß is expressed in neuronal populations, especially hippocampus and cortex. Dual labeling studies demonstrate that PI5P4Kα co-localizes with the oligodendrocyte marker, Olig2, whereas PI5P4Kß co-localizes with the neuronal marker, NeuN. Immunohistochemical subcellular distribution studies demonstrate that PI5P4Kα and PI5P4Kß are expressed in the early endosome system. Ultrastructural analysis demonstrates that both kinases are contained in axon terminals and dendritic spines adjacent to the synaptic membrane, which support a potential role in synaptic transmission. Immunohistochemical analysis of macaque and human brain tissue demonstrate a conserved pattern for PI5P4Kα and PI5P4Kß. These results highlight the diverse cell-autonomous expression of PI5P4Kα and PI5P4Kß and support further exploration into their role in synaptic function in the brain.

## Introduction

The phosphoinositide kinases, along with their corresponding phosphatases and phospholipases, regulate key functions of the phosphoinositide family of lipids. These functions include proliferation and migration, metabolic adaptation to growth acceleration, and survival in the setting of genotoxic stress (Toker, 2002). The type 2 phosphatidylinositol-5-phosphate 4-kinases (PI5P4Ks), composed of a family of 3 isoforms (α, ß and *γ*), catalyze the formation of phosphatidylinositol 4,5-bisphosphate (PI-4,5-P_2_) by phosphorylating phosphatidylinositol 5-phosphate (PI-5-P) at the 4 position of the inositol ring. These kinases play a role in a variety of disease states, including cancer, insulin signaling, and oxidative stress (Carricaburu et al., 2003; Jones et al., 2006; Keune et al., 2012; Emerling et al., 2013; Jones et al., 2013; Jude et al., 2015; Sharma et al., 2019; Wang et al., 2019).

The PI5P4K family of lipid kinases is currently thought to contribute predominantly to intracellular PI-4,5-P_2_ formation to regulate dynamics of intracellular signaling pathways. Several studies have now demonstrated that the PI5P4Ks mediate autophagy, particularly during periods of nutrient stress, by promoting autophagosome-lysosome fusion (Emerling et al., 2013; Mackey et al., 2014; Lundquist et al., 2018). These kinases also regulate cellular growth pathways, including the mammalian target of rapamycin (mTOR) and phosphatidylinositol 3-kinase (PI3K) pathways (Carricaburu et al., 2003; Gupta et al., 2013). Furthermore, they regulate early endosomal homeostasis during clathrin-mediated endocytosis, a key step in synaptic vesicle trafficking (Kamalesh et al., 2017). Of particular interest, these enzymes also play a major role in suppressing the activity of the type 1 phosphatidylinositol-4-phosphate 5-kinases through direct binding to these enzymes (Wang et al., 2019). Although the expression and function of these kinases have been explored in oncogenic signaling pathways and in settings of nutrient stress, few studies have examined their role in the brain.

PI-4,5-P_2_ is a critical mediator of synaptic function in the brain through its role in synaptic vesicle docking and fusion, leading to successful synaptic vesicle release (Honigmann et al., 2013). Dysfunction in PI-4,5-P_2_ production is associated with enhanced synaptic depression, a reduced readily releasable pool of vesicles, delayed endocytosis, and reduced recycling (Di Paolo et al., 2004). In the brain, most PI-4,5-P_2_ is created through phosphorylation of PI-4-P by the type 1 PIP kinase, PI4P5Kγ (Volpicelli-Daley et al., 2010). However, this work did not investigate the expression or localization of the PI5P4Ks in the brain.

Prior studies have demonstrated that the PI5P4Ks are highly expressed in the mouse brain as compared to other organs. PI5P4Kß localizes to the ventricular zone of the developing mouse brain, which may suggest its involvement in the stem cell niche in normal brain (Akiba et al., 2002). PI5P4Kγ, like PI4P5Kγ, is expressed in neurons and localizes to intracellular vesicles (Clarke et al., 2009). PI5P4Kα is expressed in glial cells and causes dysregulation of the glutamate transporter, excitatory amino acid transporter 3 (EAAT3) (Fedorenko et al., 2009), expressed in neurons, glial cells, and glioma. Dysregulation of the EAAT3 system has been associated with numerous neurodegenerative diseases, epilepsy, and schizophrenia (Danbolt, 2001), with large genome-wide association studies linking PIP4K2A polymorphisms to increased inherited risk of schizophrenia (Schwab et al., 2006; He et al., 2007; Rethelyi et al., 2010; Thiselton et al., 2010). While germline deletion of individual pip4k2a, pip4k2b or pip4k2c genes results in viable mice with relatively subtle phenotypes, germline deletion of both pip4k2a and pip4k2b results in neonatal lethality and deletion of pip4k2b and pip4k2c results in early embryonic lethality (Emerling et al., 2013). These results indicate that these genes have some essential but redundant functions. Though the distribution and localization of PI5P4Kγ has been well characterized in the brain, there have not been any studies examining the specific localization of PI5P4Kα and PI5P4Kß in the brain and whether this localization is conserved in mice and primates.

Given that the PI5P4Ks are highly expressed in the brain but without known expression patterns of cell type specificity, we first examined the regional localization of PI5P4Kα and PI5P4Kß in the mouse brain, using immunohistochemistry light and electron microscopy and Western blot. We then studied the localization of the PI5P4Ks in macaque and human brain tissue. This is the first comprehensive analysis of the expression and localization of PI5P4Kα and PI5P4Kß in the brain. The differential localization of these kinases within the brain may shed light on the cell-autonomous function of these kinases in the normal brain and in brain disease states.

## Materials and Methods

### Animal and human studies

A total of 9 neonatal and 63 adult mice were used in this study. Both sexes were used. pip4k2a-/- and pip4k2b-/- mice were bred on a C57BL/6 background as previously described (Lamia et al., 2004; Emerling et al., 2013). Mice were provided ad libitum food (5LA1 diet) and water in a room with a 12:12 light/dark cycle (lights on at 0600). One adult male macaque was used in this study and was maintained as previously described (Schmid et al., 2014). All mouse and macaque studies were approved by the Institutional Animal Care and Use Committee and Weill Cornell Medicine and were compliant with the 2011 8^th^ Edition of the NIH guidelines for the Care and Use of Laboratory Animals. De-identified normal post-mortem brain tissue from a male patient was obtained from the Department of Pathology at New York Presbyterian Hospital. All human studies were approved by the Institutional Review Board at Weill Cornell Medicine.

### Cell lines

All cells were incubated in a 37°C humidified incubator with 5% CO_2_. The human glioblastoma cell line, U-87MG, was obtained from American Type Culture Collection (ATCC) and was grown in DMEM containing 10% fetal bovine serum.

### Vectors and lentivirus preparation

mCherry-PIP4K2A and mCherry-PIP4K2B fusion constructs were cloned into the pLenti-CMV Puro vector. Lentivirus (LV) was prepared in 293T cells by co-transfecting mCherry-PIP4K2A and mCherry-PIP4K2B along with Pax2 and VSVG constructs. Supernatants from 293T cells were harvested after 48 hours and concentrated using LentiX concentrate (Takara Bio).

### Preparation of mouse cortical neurons and oligodendrocytes

Mouse cortical neurons were prepared as previously described(Darbinyan et al., 2013). Briefly, mouse cortices were dissected in ice-cold Hibernate E medium (Gibco) and incubated with Tryp-LE Express (Gibco) for 10 minutes with DNase I (Sigma) The cells were centrifuged, and cell pellets were resuspended in neural stem cell medium (NSCM), consisting of neurobasal medium (Gibco), N2 supplement (Gibco), B27 supplement without Vitamin A (Gibco), Glutamax (Gibco), human fibroblast growth factor (Life Technologies), human epidermal growth factor (Life Technologies), human insulin-like growth factor (Life Technologies), penicillin/streptomycin (Gibco), and gentamycin (Sigma), before being triturated with fire-polished glass pipettes. 100,000 dissociated neurospheres were plated into 4-well chamber slides and differentiated into cortical neurons through the addition of neuronal medium, consisting of neurobasal medium, B27 supplement without Vitamin A, Glutamax, human recombinant brain-derived neurotrophic factor (Life Technologies), penicillin/streptomycin, and gentamycin. After 7 DIV, 25 ul of mCherry-PIP4K2A or mCherry-PIP4K2B LV supernatant was added to neurons in chamber slides. Neurons were incubated for 2 days before performing immunofluorescence.

### Antibodies

The following commercial antibodies, which have been extensively validated elsewhere, were used: PI5P4Kα (1:500, Santa Cruz sc-100406), PI5P4Kα (1:500, Abgent AW5494), PI5P4Kß (1:500, CST 9694S), ß-actin (1:5000, Abcam ab6276), GAPDH (1:500, CST), Rab4 (1:200, Abcam, ab13252), Rab5 (1:200, CST 3547S), Rab7 (1:200, CST 9367S), EEA1 (1:200, CST 3288T). Antibody information is displayed in Table 1.

**Table 1.**
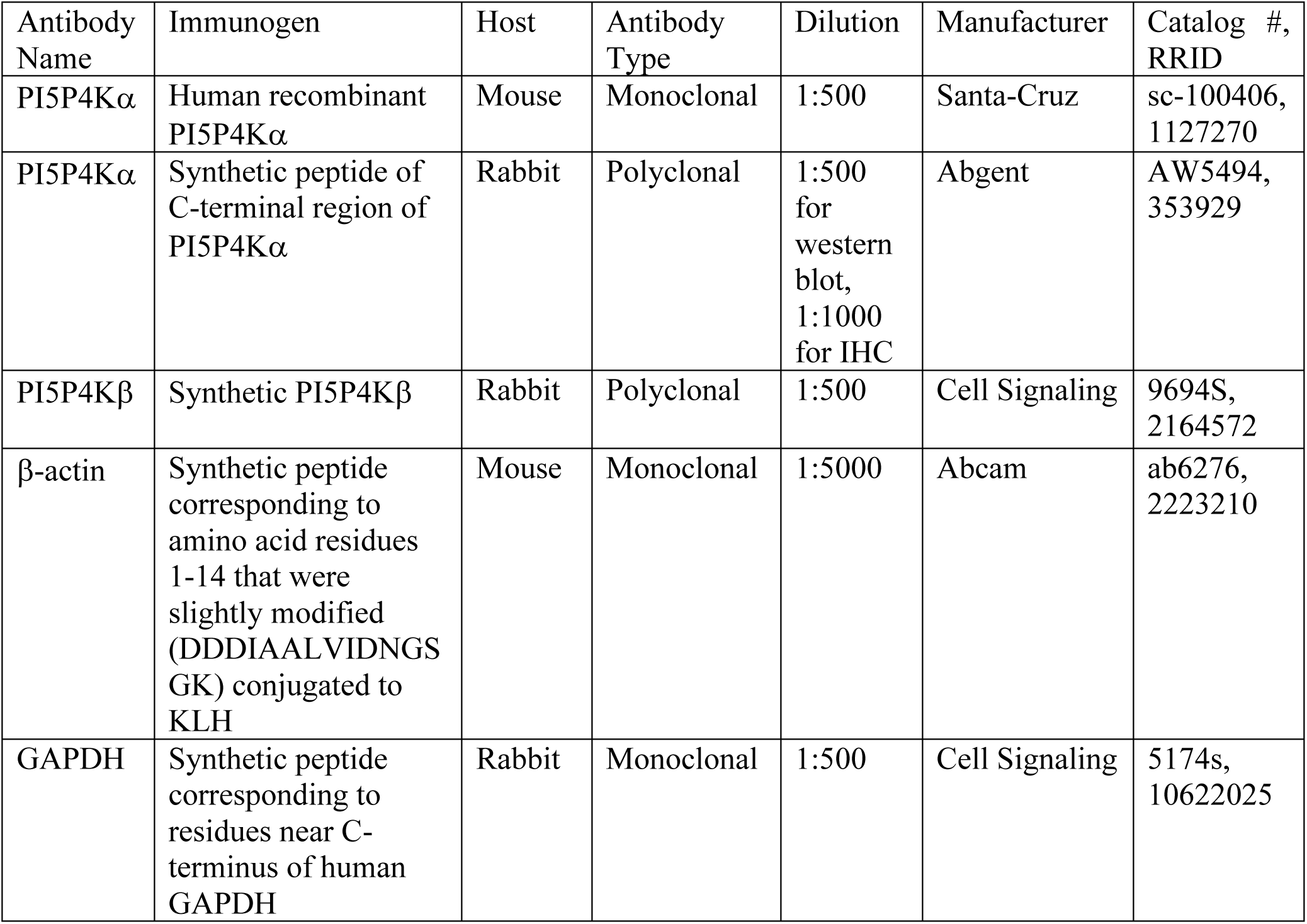
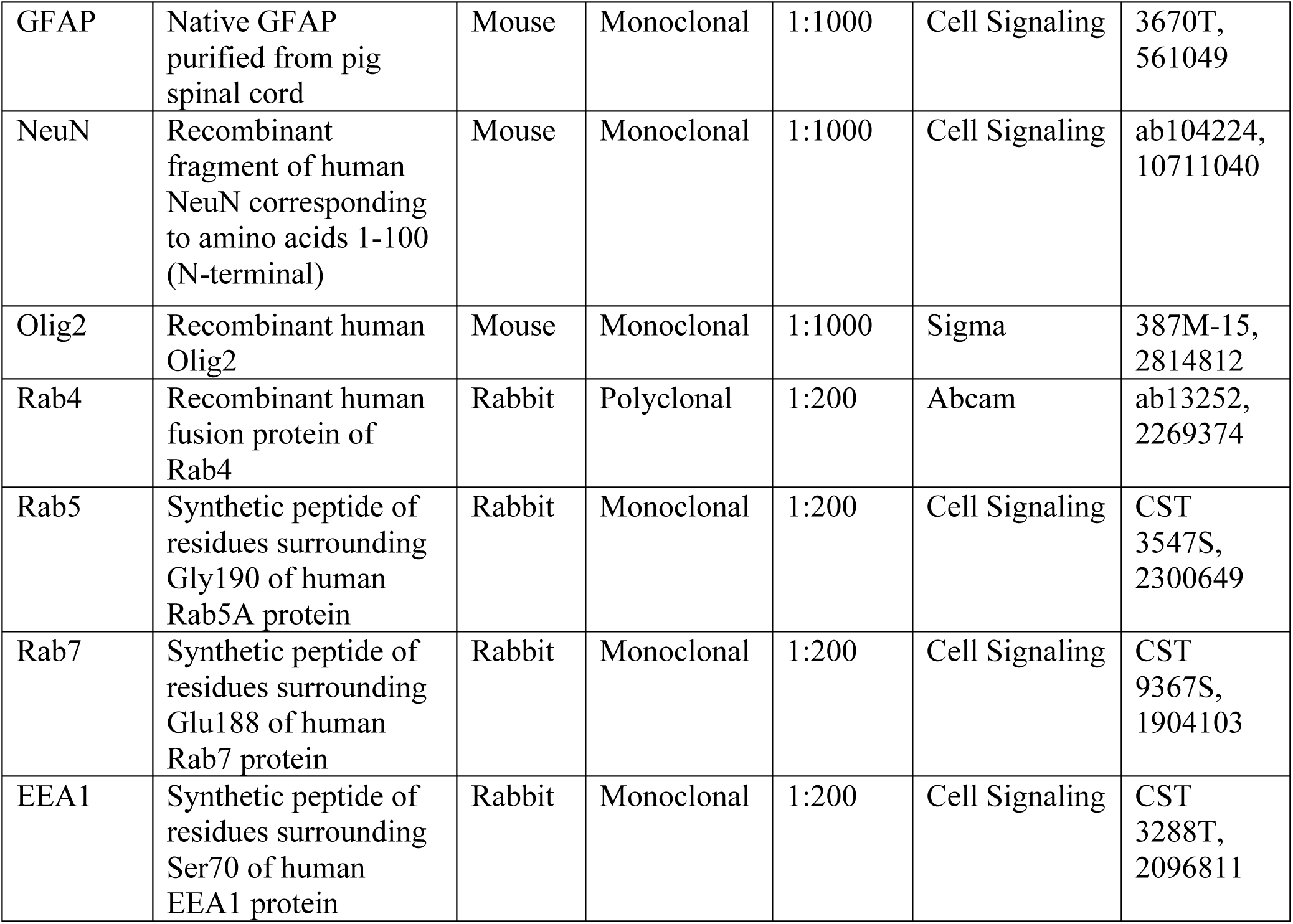
Antibody Characteristics

Specificity of PI5P4Kα antibody from Abgent and Santa-Cruz and PI5P4Kβ from Cell Signaling was tested by western blot in lysates of wild-type or pip4k2a^-/-^ or pip4k2b^-/-^ brains. Specificity of PI5P4Kα from Abgent and PI5P4Kβ antibody from Cell Signaling was tested by immunohistochemistry and immunofluorescence in brains of wild-type tissue as compared to brains of pip4k2a and pip4k2b knock-out mice, respectively.

### Western blot

Whole-cell lysates were prepared from whole mouse brains using CST buffer (Cell Signal Technology) containing protease inhibitor (Sigma), and were rotated at 4°C for 30 minutes before DNA was pelleted. Total protein (30 μg) was loaded on 4-12% Tris-Glycine gels (Thermo-Fisher) for SDS-PAGE. Blots were incubated with the following primary antibodies: PI5P4Kα (1:500, Santa Cruz sc-100406), PI5P4Kα (1:500, Abgent AW5494), PI5P4Kß (1:500, CST 9694S), ß-actin (1:5000, Abcam), and GAPDH (1:500, CST). Blots were then incubated with either mouse or rabbit secondary antibodies as appropriate, and developed using Dura or Femto enhanced chemiluminescence (Pierce).

### Immunocytochemistry

Cortical neurons or oligodendrocytes were cultured in chamber slides and then fixed in 4% paraformaldehyde (PFA) in 1x phosphate-buffered saline (PBS) for 15 minutes. Cells were washed for 5 minutes in PBS and then incubated in blocking buffer containing 5% normal donkey serum and 0.1% bovine serum albumin (BSA) for 1 hour. Cells were washed for 5 minutes in PBS and then incubated overnight at 4°C with the following primary antibodies: Rab4, Rab5 (1:200, CST 3547S), Rab7 (1:200, CST 9367S), EEA1 (1:200, CST 3288T), Lamin A/C (1:200, CST 4777T), Clathrin (1:200, CST, Synaptophysin (1:200, Abcam ab32127), Synaptotagmin (1:200, Abcam ab13259, Synaptobrevin (1:200, Synaptic Systems 104211). The following day, cells were washed 3 times in PBS for 5 minutes before being incubated with AlexaFluor 488 or AlexaFluor 568 (Thermo Fisher) rabbit or mouse secondary antibodies for 1 hour at room temperature. Cells were then mounted with Prolong Gold Antifade Mountant containing DAPI (Thermo Fisher) and coverslipped.

### Immunohistochemistry and immunofluorescence

For immunohistochemistry, mice were injected with sodium pentobarbital (150 mg/kg, i.p.) and perfused transcardially with 2% heparin-saline followed by 30 ml 3.75% acrolein and 2% PFA in 0.1 M phosphate buffer (PB; pH 7.4). The brains were removed from the skull, post-fixed in 1.9% acrolein and 2% PFA in PB for 30 minutes, and then placed in PB. Coronal sections through the brains were cut (40-µm thick) using a Vibratome (Leica Microsystems) and stored in cryoprotectant at –20°C until use (Milner et al., 2011).

For embryonic brain analysis, cervical dislocation was performed on pregnant mothers containing fetuses at E15.5 or E18.5. Mouse fetuses were removed, and brains were extracted from the skull. Whole brains were submerged in 4% PFA in PBS at room temperature for 2 days and then for 1 day at 4°C. PFA was replaced with PBS containing sodium azide and stored at 4°C until ready to use. Post-natal mice underwent cervical dislocation, and brains were removed, submerged in 4% PFA in PBS as above, and stored in PBS containing sodium azide until ready to use. For embryonic and post-natal brains used for determining PI5P4Kα and PI5P4Kß localization across development, whole brains were prepared in sheets embedded in matrix (NeuroScience Associates Labs, Knoxville, TN). Free-floating sheets containing brains were then used for subsequent experiments.

For immunohistochemistry and immunofluorescence, all brain sections were incubated in blocking buffer (5% normal donkey serum (Jackson ImmunoResearch) and 0.3% Triton-X 100 (Sigma) in PBS, pH 7.4 and avidin/biotin blocking kit (Vector Laboratories, SP-2001) at room temperature, and then incubated with primary antibodies in blocking buffer for 24 hours at room temperature followed by 48 hours at 4°C. The following primary antibodies were used: rabbit anti-PI5P4Kα (1:500, Abgent AW5494), rabbit anti-PI5P4Kß (1:1000, CST 9694S), mouse anti-Olig2 (1:1000, Sigma 387M-15), mouse anti-GFAP (1:1000, CST 3670T), mouse anti-NeuN (1:1000, Abcam ab104224). For immunohistochemistry, the secondary antibodies used were biotinylated donkey anti–rabbit IgG (1:400, Jackson ImmunoResearch, catalog 713-065-147). For immunofluorescence, the secondary antibodies used were donkey anti-rabbit AlexaFluor 488 and donkey anti-mouse AlexaFluor 568 (Thermo Fisher). Sections were coverslipped with Prolong Gold Antifade Mountant containing DAPI (Thermo Fisher).

Brain regions in embryonic, post-natal, and adult mice were identified using developmental (Paxinos, 1991; Altman and Bayer, 1995) and adult brain atlases (Hof et al., 2000). Three mice at each pre- and post-natal age were analyzed. For light microscopy, images were acquired on a Zeiss Axio Observer.Z1 inverted microscope and captured using the same exposure time/channel/experiment between different genotypes by AxioCam MRC camera. For immunofluorescence, images were acquired on a Zeiss LSM 880 Laser Scanning Confocal Microscope equipped with ZEN software (Zeiss).

### Electron microscopy

Sections were processed for electron microscopy as previously described (Milner et al., 2011). Briefly, 4-8-week-old WT, pip4k2a-/- mice, and pip4k2b-/- mice were overdosed with sodium pentobarbital (150 mg/kg, i.p.) and perfused transcardially with 2% heparin-saline followed by 30 ml, 3.75% acrolein, and 2% PFA in PB. The brains were removed from the skull, post-fixed in 1.9% acrolein and 2% PFA in PB for 30 minutes, and then placed in PB. Coronal sections through the brains were cut (40-µm thick) using a Vibratome (Leica Microsystems) and stored in cryoprotectant at –20°C until use.

For PI5P4Kα and PI5P4Kß staining, coronal sections (n = 3 animals per genotype) were rinsed in PB to remove cryoprotectant and then incubated in 1% sodium borohydride in PB for 30 minutes to remove unbound aldehydes. Sections were washed in 8–10 changes of PB until all the gaseous bubbles disappeared and then placed in 0.1 M Tris-buffered saline (TS), pH 7.6. Sections were then incubated on a shaker sequentially in: (i) 0.5% BSA in TS (30 min); (ii) primary antibodies (rabbit anti-PI5P4Kα (1:500, Abgent) and rabbit anti–PI5P4Kß (1:1000, CST 9594) in 0.1% BSA in TS for 1 day at room temperature (∼23°C), followed by 2 days at 4°C (iii) 1:400 of biotinylated anti-goat IgG (Jackson ImmunoResearch, catalog 705-065-147), 30 minutes; (iv) 1:100 peroxidase-avidin complex (Vectastain Elite Kit, Vector Laboratories, catalog PK-6100), 30 minutes; and (v) DAB (Sigma-Aldrich) and H_2_O_2_ in TS, 3 minutes. All incubations were separated by 2-3 washes in TS.

For immunogold detection of PI5P4Kα and PI5P4Kß, DAB-reacted sections were rinsed in 0.01 M PBS (pH 7.4), incubated in blocking buffer (0.8% BSA, 0.2% gelatin, 0.02% BSA in 0.02 M PBS) for 30 minutes, and placed overnight in a 1:50 dilution of donkey anti–rabbit IgG with bound 10-nm colloidal gold (Electron Microscopy Sciences, catalog 25704) diluted in blocking buffer. The gold particles were fixed to the tissue in 2% glutaraldehyde in 0.01 M PBS and rinsed in PBS followed by 0.2 M sodium citrate buffer (pH 7.4). The bound silver-gold particles were enhanced using a Silver IntenSE M kit (catalog RPN491; GE Healthcare) for 7 minutes.

Sections were rinsed in 0.1 M PB and then post-fixed in 2% osmium tetroxide in PB for 1 hour, dehydrated, embedded with Epon 812 (Electron Microscopy Sciences) between 2 sheets of Aclar plastic, and left to cure at 60°C overnight. Sections were selected, mounted on EMBed chucks (Electron Microscopy Sciences) and trimmed to 1–1.5 mm–wide trapezoids. Ultrathin sections (∼65 nm thick) within 0.1–0.2 µm to the tissue-plastic interface were cut on a Leica Ultracut ultratome, collected into copper mesh grids, and counterstained with uranyl acetate and Reynold’s lead citrate. Sections were viewed and photographed using a FEI Tecnai Biotwin electron microscope equipped with a digital camera (Advanced Microscopy Techniques, software version 3.2).

### Ultrastructural image analysis

The data analysis procedure is similar to previously described (Marques-Lopes et al., 2015). The dendritic profiles contained regular microtubular arrays and were usually postsynaptic to axon terminal profiles (Peters et al., 1991). Dendritic spines were small (about 0.1–0.2 µm in diameter), abutted terminals, and sometimes emanated from dendritic shafts (Peters et al., 1991). Immunoperoxidase labeling for PI5P4Kα and PI5P4Kß was distinguished as an electron-dense reaction product precipitate. For quantification, all labeled axon terminals and dendritic spines from the CA1 region of hippocampus from either PI5P4Kα or PI5P4Kß-stained brain sections were counted from two grid squares (6050 μm^2^) from 3 mice per condition.

#### Primate tissue

A male Macaque was deeply anesthetized with ketamine 15 mg/kg IM potentiated by xylazine 2 mg/kg IM (Rompun, Haver) and transcardially perfused with 4% PFA in PB. Coronal sections (40 μm thick) were cut on a vibratome and stored in cryoprotectant at –20°C until use.

### Human tissue samples and IHC

Post-mortem human brain tissue from the temporal lobe (containing the hippocampus) was post-fixed in acrolein in PBS and sectioned (40 μm thick) on a vibratome and stored in cryoprotectant at –20°C until use. IHC was performed using the primary antibodies, PI5P4Kα antibody (1:1000, Abgent AW5494) and PI5P4Kß antibody (1:500, CST 9694S), followed by secondary antibody incubation with biotinylated donkey anti-rabbit secondary antibodies and then DAB incubation. Sections were mounted on gelatin coated slides, dehydrated and coverslipped with Permount (Fisher). Slide images were captured with an Observer.Z1 (Carl Zeiss), and digital images were acquired with an AxioCam MRc camera and AxioVision 4.8.

#### Statistics

All of the data were analyzed with the program GraphPad Prism version 7.0 (GraphPad Software). A two-tailed student’s t-test was used to analyze significance between groups in electron microscopy experiments.

## Results

### Antibody Characterization

Previous studies have localized PI5P4K*γ* protein and PIP4K mRNA in mouse brain (Clarke et al., 2009), but no studies have examined localization of PI5P4Kα and PI5P4Kβ protein in the brain of any species. We tested the specificity of PI5P4Kα and PI5P4Kβ antibody in wild-type and pip4k2a^-/-^ and pip4k2b^-/-^ knock-out mice by Western blot and immunohistochemistry using two commercially available antibodies (Fig. 1A). We also demonstrate specificity of PI5P4Kα and PI5P4Kβ antibodies in wild-type and respective knockout mouse brain tissue by immunohistochemistry (Fig. 1B) and immunofluorescence (Fig. 1C). PI5P4Kα immunolabeling was found throughout the corpus callosum in wild-type, but not knockout, sections (Fig. 1B, C). Likewise, PI5P4Kβ immunolabeling was found in the hippocampal pyramidal cell layer in wild-type, but not knockout mice (Fig. 1B, C).

**Figure 1.**
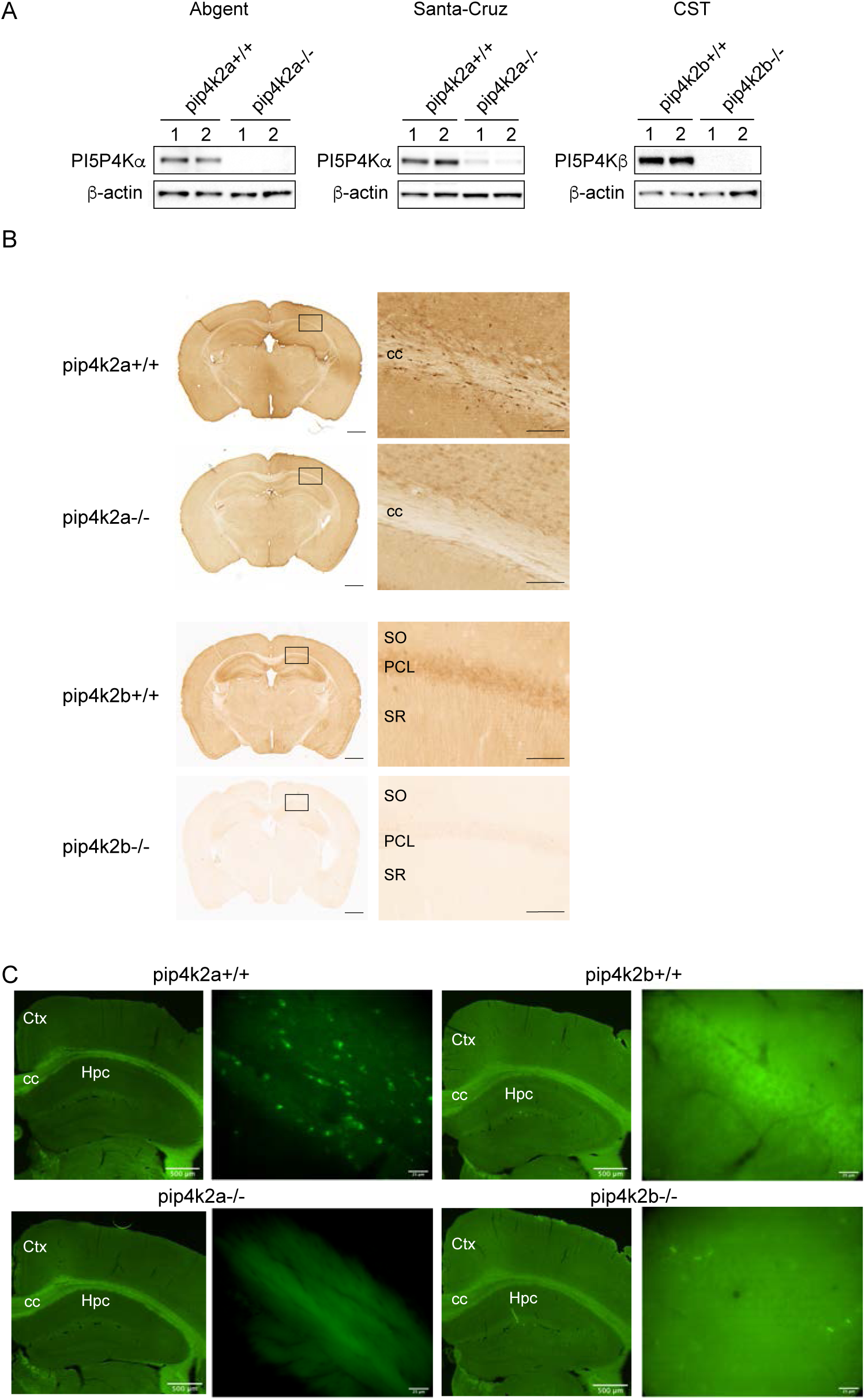
Confirmation of antibody specificity for PI5P4Kα and PI5P4Kβ in mouse brain A. Western blot analysis of whole brain lysates in wild-type and pip4k2a knockout mouse brain shows specificity of 2 antibodies (Abgent and Santa-Cruz) for PI5P4Kα, and specificity of Cell Signal (CST9594S) for PI5P4Kβ in wild-type compared to knockout brains. B. Immunohistochemical expression for PI5P4Kα (left) and PI5P4Kβ (right) in wild-type and knockout brains shows positive signal in wild-type brains and lack of staining in pip4k2a and pip4k2b knockout brains. Scale bars for immunohistochemical images are 1 mm for low-magnification images on left and 125 μm for high-magnification images on right. C. Immunofluorescence for PI5P4Kα (left) and PI5P4Kβ (right) in wild-type and knockout brains shows positive PI5P4Kα signal in corpus callosum and positive PI5P4Kβ signal in hippocampus in wild-type brains and lack of staining in pip4k2a and pip4k2b knockout brains. Scale bar length is indicated for each bar. CC, corpus callosum; PCL, pyramidal cell layer; SO, stratum oriens; SR, stratum radiatum; Hpc, hippocampus; Ctx, cortex

### PI5P4Kα and PI5P4Kß are differentially expressed in mouse brain during development

To characterize the expression of PI5P4Kα and PI5P4Kβ in mouse brain across development, we performed Western blot of whole brain lysates from mouse brains at 3 embryonic time-points and 12 post-natal time-points. We found that expression of PI5P4Kα increased in post-natal stages, around P11, further increasing up until p30 (Fig. 2A). On the other hand, PI5P4Kβ expression remained constant from embryonic day 12.5 (E12.5) to post-natal day 30 (p30). These data correlate with findings from the BrainRNASeq database, a transcriptome and splicing database of glia, neurons, and vascular cells of the cerebral cortex (Zhang et al., 2014). This database shows that PIP4K2A expression increases from early post-natal development at p7 up until 9.5 months and 2 years of age (Fig. 2B) and that PIP4K2B expression is highest at early post-natal development at p7 and decreases thereafter (Fig. 2B).

**Figure 2.**
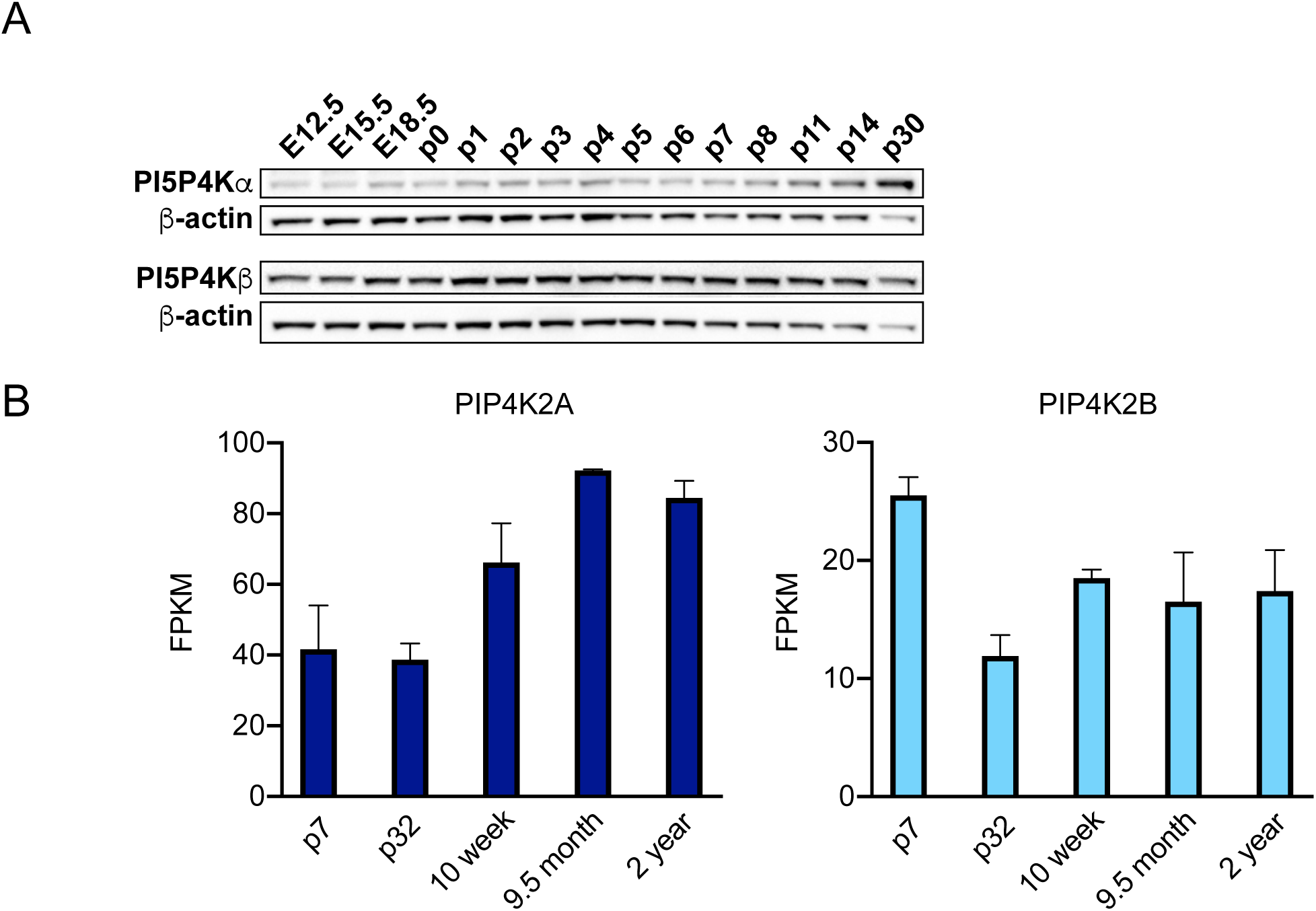
Developmental expression of PI5P4Kα and PI5P4Kβ in the mouse brain A. Western blot analysis of PI5P4Kα and PI5P4Kβ in whole brain lysates at the indicated developmental timepoints. B. RNAseq analysis of PI5P4Kα and PI5P4Kβ expression from the BrainRNAseq database at the indicated developmental timepoints.

By immunohistochemistry, we evaluated localization of PI5P4Kα and PI5P4Kβ across development. We found that PI5P4Kα-immunolabeling was contained in white matter, particularly in corpus callosum, beginning around p14 (Fig. 3). We also found PI5P4K*α−*labeled cells in white matter tracks in the cerebellum, including the arbor vitae, beginning at E18.5 and becoming more prominent by p28 (Fig. 3). PI5P4Kα expression was absent in the molecular and granular layers of the cerebellum at all time-points.

**Figure 3.**
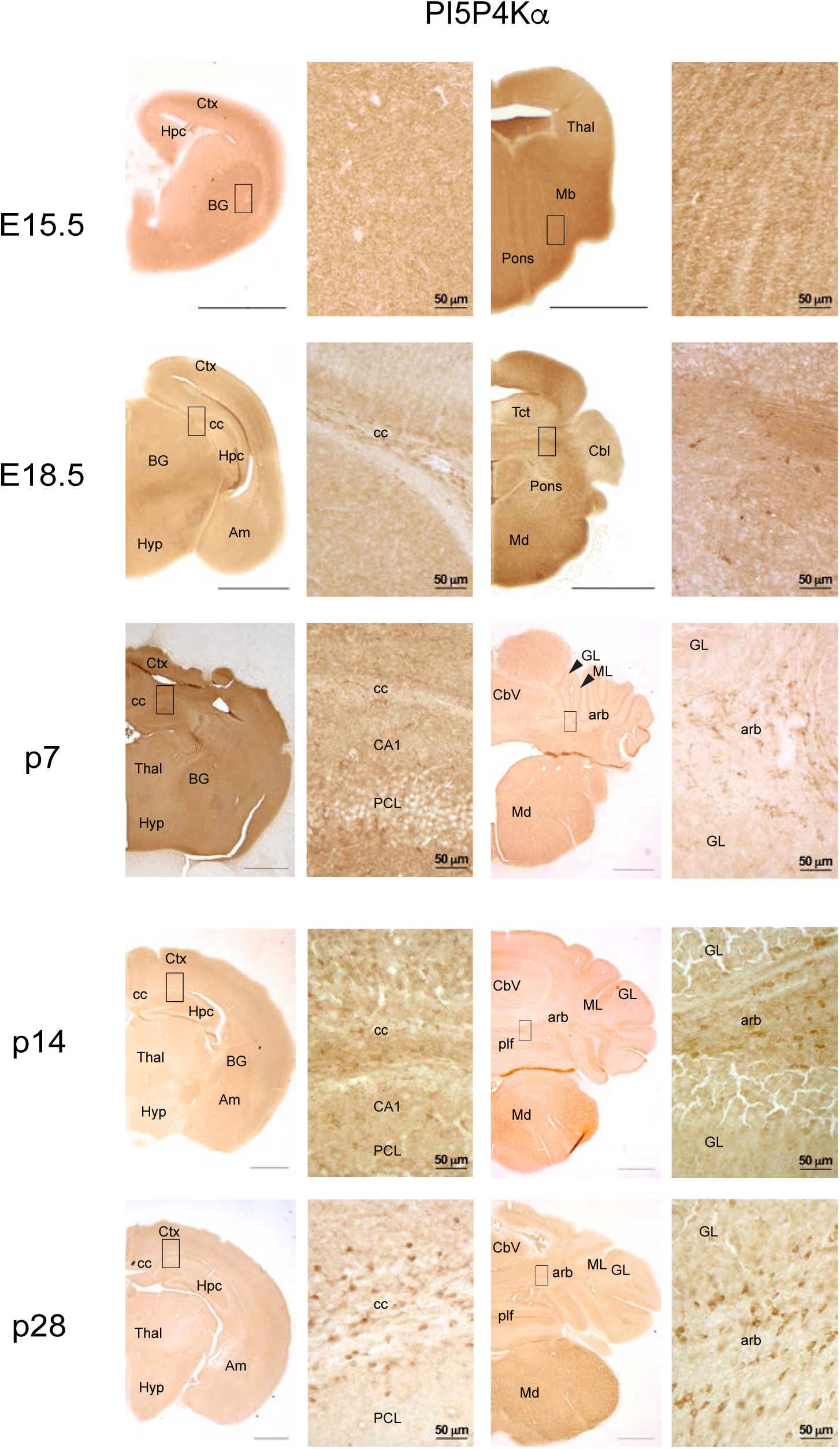
Immunohistochemical analysis for PI5P4Kα at the indicated developmental timepoints. Scale bars for low-magnification images = 150 μm, scale bars for high-magnification images are indicated for each bar; Ctx, cortex; Hpc, hippocampus; BG, basal ganglia; Thal, thalamus; Mb, midbrain; Hyp, hypothalamus; Am, amygdala; Tct, tectum; Cbl, cerebellum; Md, medulla; CC, corpus callosum; PCL, pyramidal cell layer; Cbv, cerebellar vermis; GL, granule cell layer of the cerebellum; ML, molecular cell layer of the cerebellum; plf, posterolateral fissure; arb, arbor vitae; arrowheads identify relevant named structures

PI5P4Kβ-immunolabeling was evident early in development at E15.5, especially in cortex, basal ganglia, and hippocampus. In the E15.5 cortex, labeled cells were densest in the cortical plate. However, by E18.5, PI5P4Kβ-positive cells were primarily found in the ventricular zone. This pattern of labeling was similar at the remaining developmental time points. At E15.5, PI5P4Kβ-labeled cells were found scattered throughout the basal ganglia with a greater concentration in the region of the lateral migratory steam. The distribution of PI5P4K*β−*labeled cells in the basal ganglia was similar at post-natal stages of development (not shown). The E15.5 hippocampus contained PI5P4Kβ-labeled cells, but the laminae were indistinct at this time point. By E18.5, cells in the pyramidal cell layer, especially CA1, were prominently labeled with PI5P4Kβ. By p7, PI5P4Kβ-immunoreactivity was distinguishable within pyramidal cells and interneurons as well as granule cells (Fig. 4). In the cerebellum, PI5P4Kβ expression was evident in the granular layer beginning at p7 and in the molecular layer beginning at p14. PI5P4Kβ expression was absent in the arbor vitae of the cerebellum at all time-points. These findings demonstrate that PI5P4Kα and PI5P4Kβ exhibit not only differential levels of expression during development but also regional diversity throughout the mouse brain.

**Figure 4.**
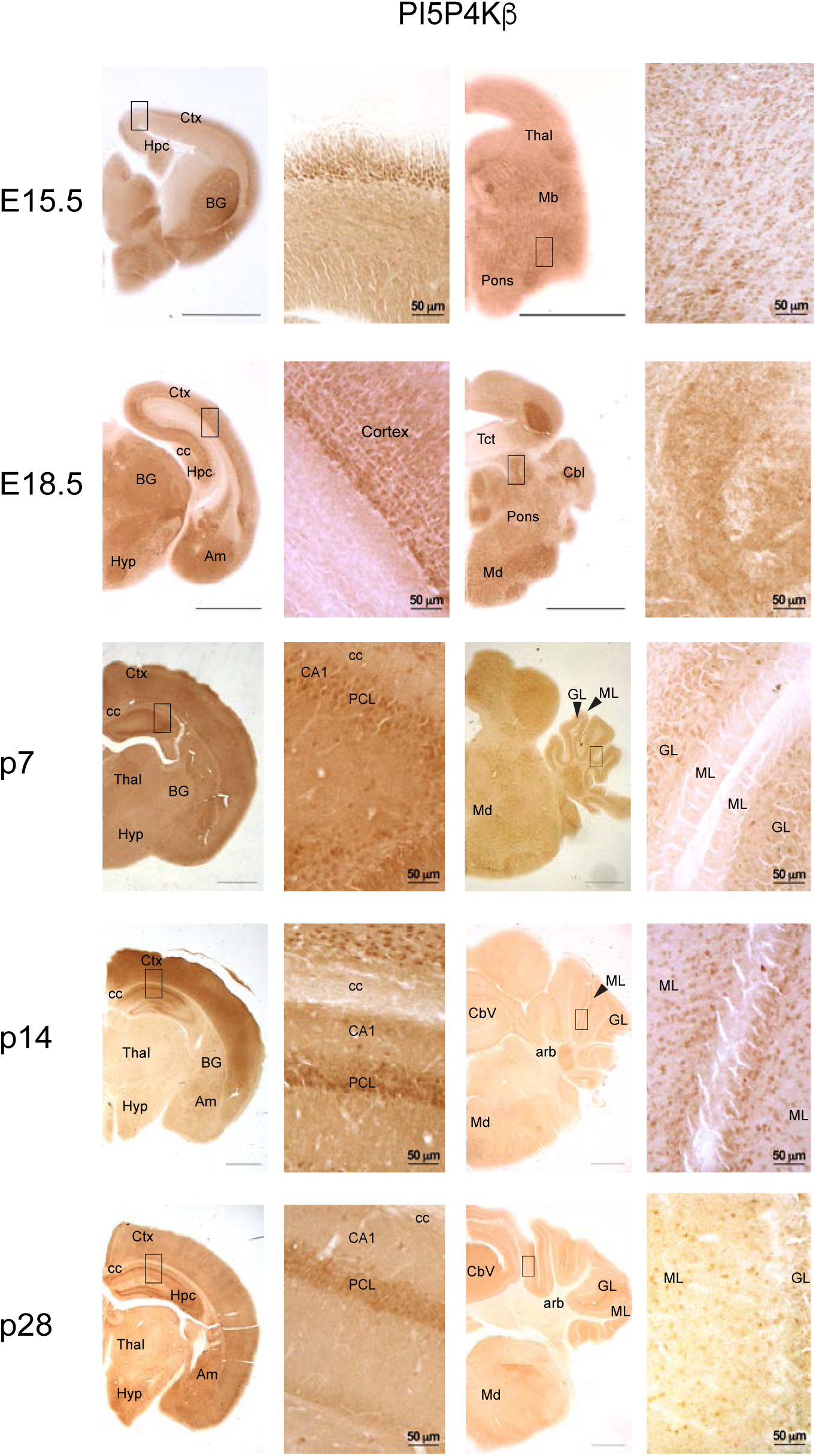
Immunohistochemical analysis for PI5P4Kβ at the indicated developmental timepoints. Scale bars for low-magnification images = 150 μm, scale bars for high-magnification images are indicated for each bar; Ctx, cortex; Hpc, hippocampus; BG, basal ganglia; Thal, thalamus; Mb, midbrain; Hyp, hypothalamus; Am, amygdala; Tct, tectum; Cbl, cerebellum; Md, medulla; CC, corpus callosum; PCL, pyramidal cell layer; Cbv, cerebellar vermis; GL, granule cell layer of the cerebellum; ML, molecular cell layer of the cerebellum; plf, posterolateral fissure; arb, arbor vitae; arrowheads identify relevant named structures

### PI5P4Kα and PI5P4Kβ are differentially expressed in the adult mouse brain

To further differentiate the regional levels of PI5P4Kα and PI5P4Kβ expression in the mouse brain, we determined the localization of PI5P4Kα and PI5P4Kβ in 9 different brain regions by dissection of each region prior to preparing whole cell lysates and subsequent Western blot. We found that PI5P4Kα and PI5P4Kβ are expressed in each region of the brain and spinal cord that we assayed (Fig. 5 A,B). However, we found that both PI5P4Kα and PI5P4Kβ expression are highest in cortex and lowest in spinal cord, with PI5P4Kβ also having relatively high levels of expression in olfactory bulb, caudate putamen, thalamus, hippocampus and midbrain.

**Figure 5.**
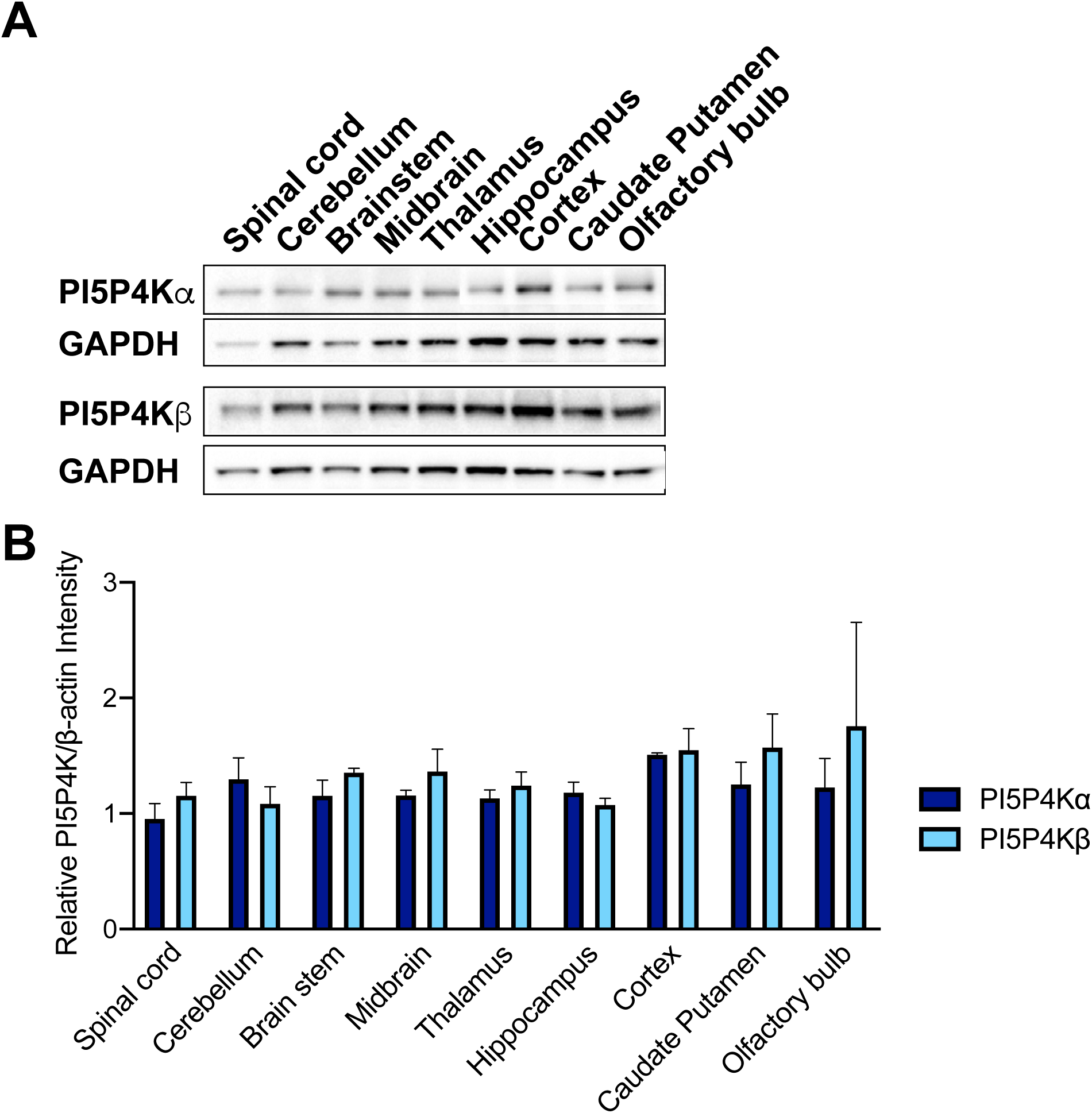
Differential expression of PI5P4Kα and PI5P4Kβ in the mouse brain by western blot analysis A. Western blot of lysates obtained from the indicated brain regions. B. B. Quantification of western blot band intensities from A (n=3 mice per group).

We next surveyed the expression of PI5P4Kα and PI5P4Kβ in the adult mouse brain through immunohistochemical analysis. PI5P4Kα expression is highest in the corpus callosum, with a scattered pattern of expression in populations of cells within layer 5 of cortex, the CA1 region of hippocampus, the ventral posteromedial nucleus of the thalamus, and the cerebellar commissure and funicular nucleus of the cerebellum (Fig. 6A and B). Conversely, strong PI5P4Kβ expression was found throughout the brain in the hippocampal pyramidal cell layer, layers 4 and 5 of cortex, ventral posteromedial nucleus of the thalamus, and laterodorsal tegmental nucleus (Fig. 6A and B). Expression of PI5P4Kβ was absent within the corpus callosum.

**Figure 6.**
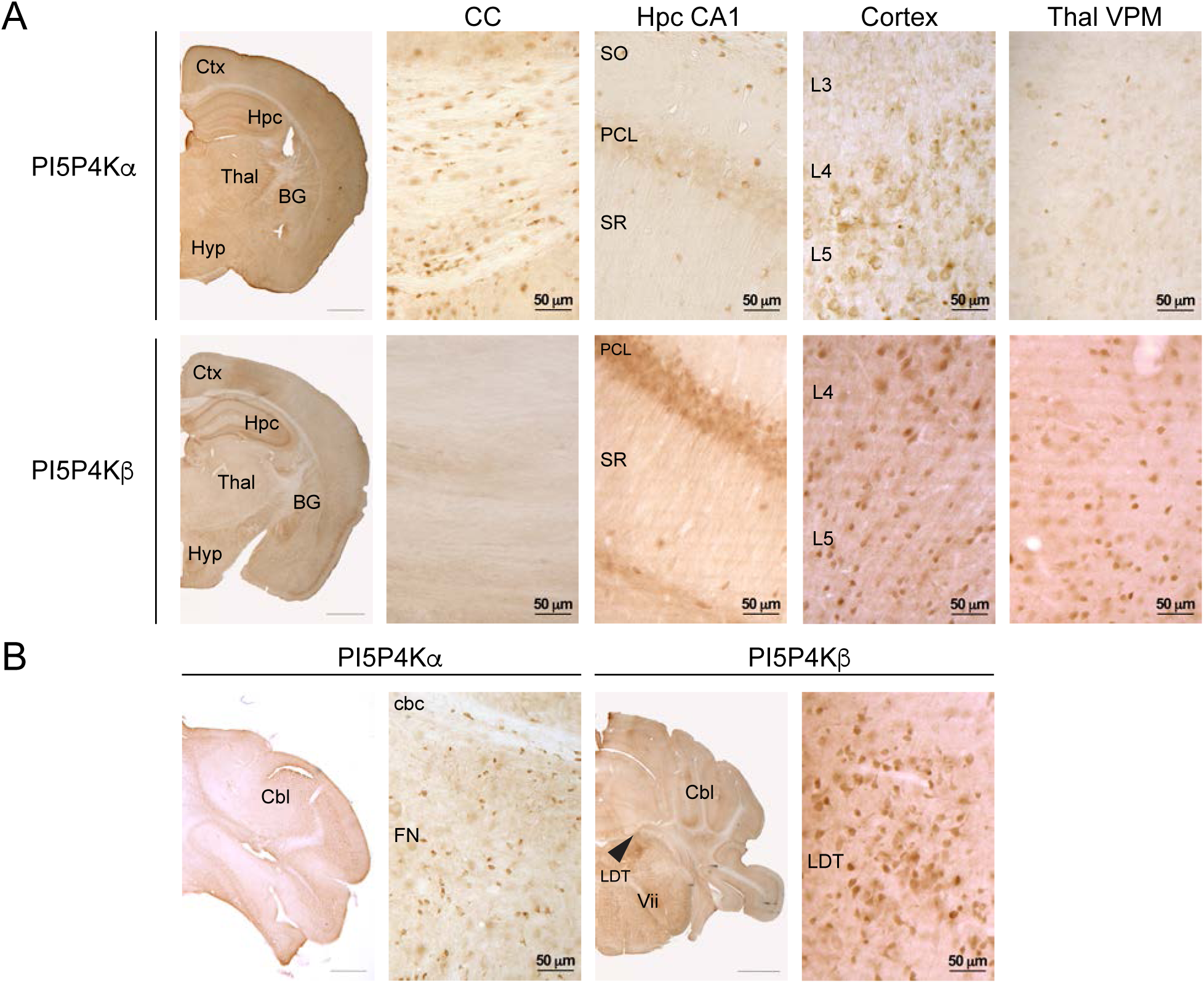
Immunohistochemical distribution of PI5P4Kα and PI5P4Kβ in the mouse brain A. At the level of rostral hippocampus (top set of panels), PI5P4Kα is expressed in oligodendrocytes throughout the corpus callosum, in neurons and oligodendrocytes in cortex and in oligodendrocytes in the midbrain. PI5P4Kα is also expressed at low levels in neurons in hippocampus. In the cerebellum (bottom set of panels), PI5P4Kα is expressed in oligodendrocytes and some neurons. B. At the level of rostral hippocampus (top set of panels), PI5P4Kβ is expressed in neurons in the hippocampal pyramidal cell layer, in cortical neurons, and in neurons in the midbrain. PI5P4Kβ expression is absent in the corpus callosum. In the bottom set of panels, PI5P4Kβ is expressed in neurons in the laterodorsal tegmental nucleus (arrowhead). Ctx, cortex; Hpc, hippocampus; BG, basal ganglia; Thal, thalamus; Hyp, hypothalamus; Cbl, cerebellum; CC, corpus callosum; PCL, pyramidal cell layer; SO, stratum oriens; SR, stratum radiatum; L3, cortical layer 3; L4, cortical layer 4, L5, cortical layer 5; cbc, cerebellar commissure; FN, funicular nucleus; LDT, laterodorsal tegmental nucleus; Vii, 7^th^ cranial nerve. Scale bars for low-magnification images = 1 mm, scale bars for high-magnification images are indicated for each bar.

### PI5P4Kα and PI5P4Kβ are expressed in different cell types in mouse brain, and this differential expression is also observed in macaque, and human brain

To determine the cellular identity of PI5P4Kα and PI5P4Kβ-expressing cells in the mouse brain, we performed immunofluorescent analysis with cell type-specific markers. Using double labeling immunofluorescence, we found that PI5P4Kα expression is restricted mainly to oligodendrocytes and some neuronal populations, whereas PI5P4Kβ expression is restricted to neurons. PI5P4Kα co-localized with the oligodendrocyte marker, Olig2, in corpus callosum but did not co-localize with the astrocyte marker, GFAP (Fig. 7A-C). PI5P4Kα was expressed at a low level in neurons within the hippocampal pyramidal cell layer. On the other hand, diffuse PI5P4Kβ immunoreactivity was visualized in NeuN-labeled neurons throughout the brain, but did not co-localize with Olig2 or GFAP (Fig. 7D-F). We performed co-localization studies in macaque and human brains to confirm our findings in the mouse brain. We found PI5P4Kα expression in oligodendrocytes in macaque and human corpus callosum and neurons in macaque and human hippocampus, and we found PI5P4Kβ expression in neurons in macaque and human hippocampus (Fig. 8A and B). These findings indicate conservation of cell type specificity for PI5P4Kα and PI5P4Kβ among mouse, macaque, and human species.

**Figure 7.**
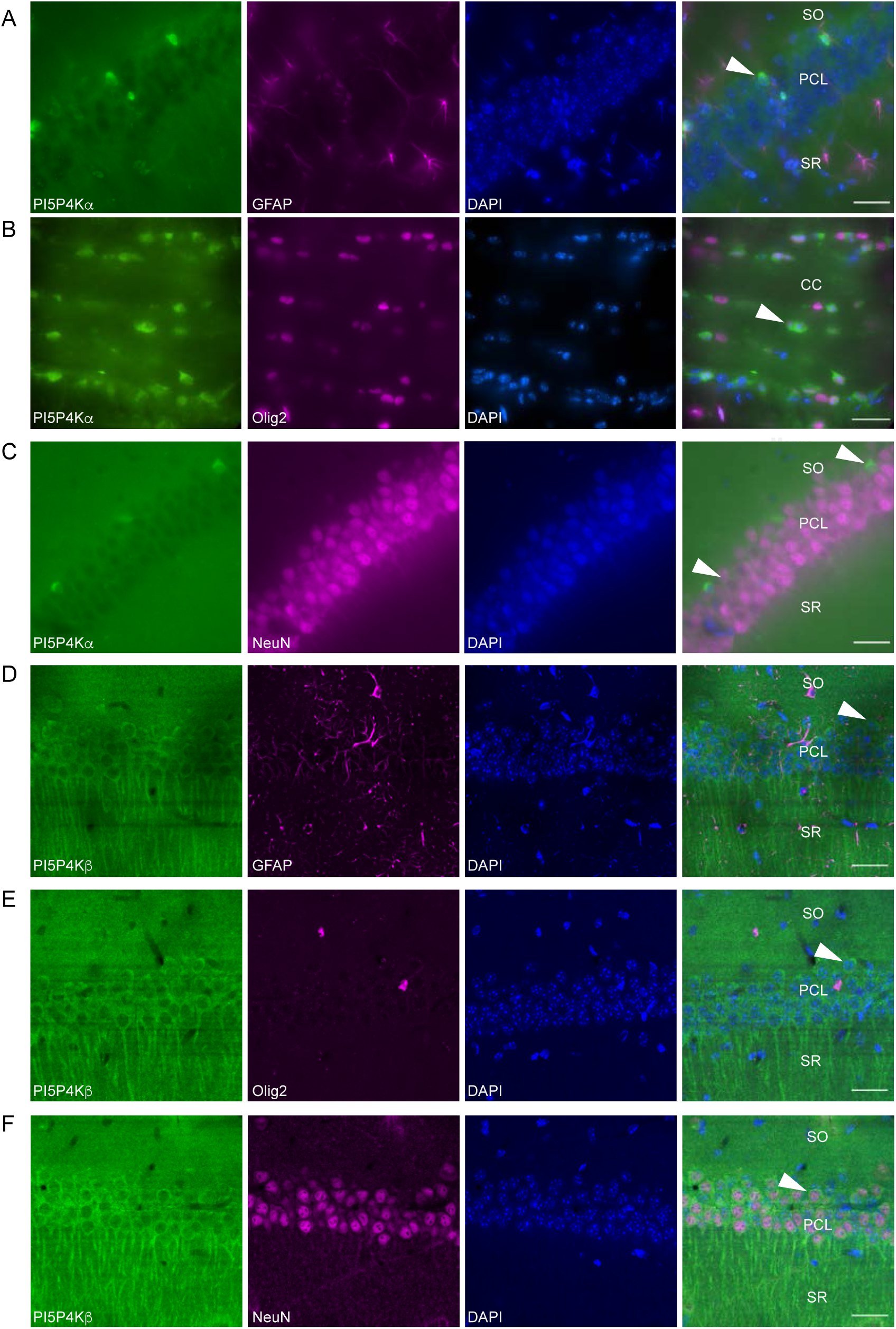
Immunofluorescent characterization of cell type expression of PI5P4Kα and PI5P4Kβ in the mouse brain PI5P4Kα does not localize with GFAP in the hippocampal pyramidal cell layer (A) but localizes with Olig2 in corpus callosum (B) and is present in some neurons within the hippocampal pyramidal cell layer (C). PI5P4Kβ localizes with NeuN (F) but not Olig2 (D) or GFAP (E) in the hippocampal pyramidal cell layer. Arrowheads identify positive staining for PI5P4Kα and PI5P4Kβ. CC, corpus callosum; PCL, pyramidal cell layer; SO, stratum oriens; SR, stratum radiatum Scale bars = 30 μm.

**Figure 8.**
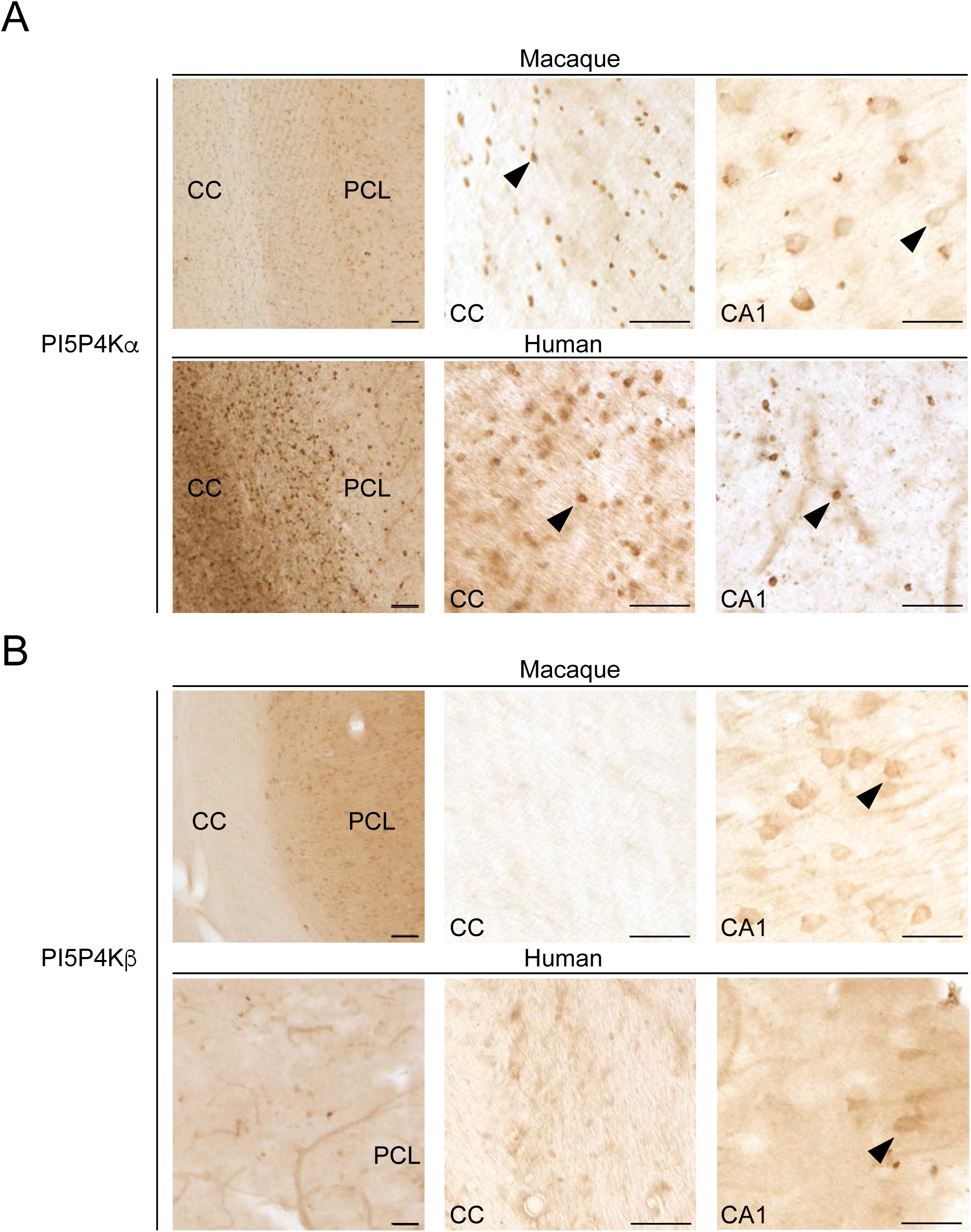
Immunohistochemical distribution of PI5P4Kα and PI5P4Kβ in the macaque and human brain A. PI5P4Kα is expressed in oligodendrocytes (arrowheads) in corpus callosum and neurons in macaque hippocampus. PI5P4Kα is expressed in oligodendrocytes in human corpus callosum and hippocampus. B. PI5P4Kβ is expressed in neurons (arrowheads) in hippocampus in macaque and human brain. Arrowheads identify positive staining for PI5P4Kα and PI5P4Kβ. CC, corpus callosum; PCL, pyramidal cell layer. Scale bars on left panel = 150 μm; scale bars on right 2 panels = 50 μm.

Data from the BrainRNASeq database are in agreement with our immunohistochemical findings of PI5P4Kα and PI5P4Kβ protein levels (Zhang et al., 2014; Zhang et al., 2016). We probed this database to determine the mRNA expression levels of PIP4K2A and PIP4K2B. In the mouse brain, PIP4K2A mRNA expression is highest in oligodendrocytes but is much lower in neurons (Fig. 9A). Interestingly, PIP4K2B mRNA is also found in microglia. Similarly, in human brain, PIP4K2A mRNA expression is highest in oligodendrocytes. For PIP4K2B, mRNA expression is highest in neurons in mouse brain, whereas in human brain, astrocytes, oligodendrocytes, and neurons express nearly similar levels of PIP4K2B mRNA (Fig. 9B). The oligodendrocyte-specific localization of PIP4K2A is also supported by a recent paper demonstrating that PIP4K2A is a marker gene for oligodendrocytes (Schirmer et al., 2019).

**Figure 9.**
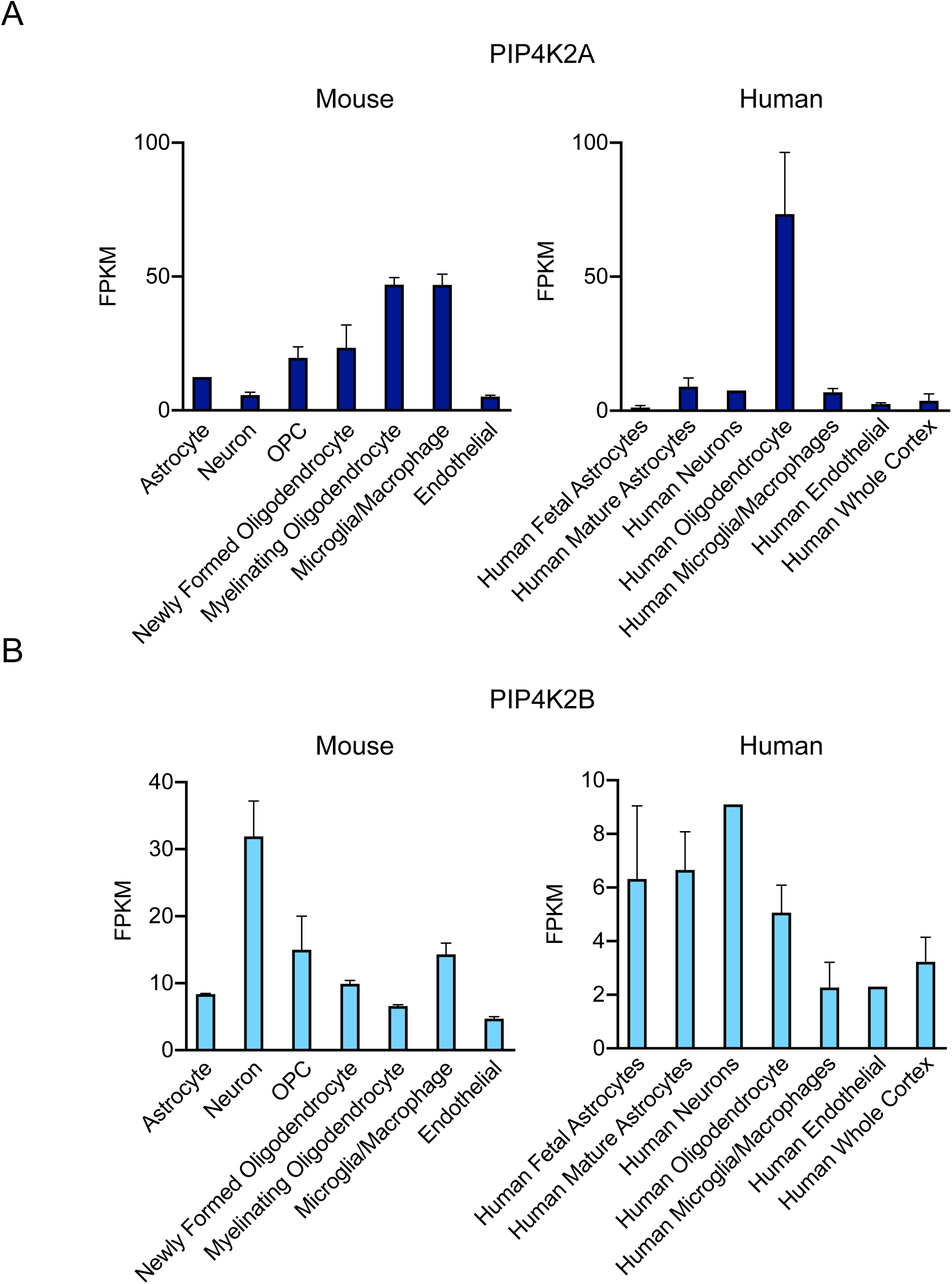
Cell type distribution of PIP4K2A and PIP4K2B throughout whole mouse brain RNAseq analysis A. Pip4K2A is expressed mostly in oligodendrocyte lineage cells in mouse and human brain, with microglia and macrophages also displaying high levels of expression. B. PIP4K2B is expressed mostly in neurons in mouse brain and in astrocytes and oligodendrocytes in human brain.

### PI5P4Kα and PI5P4Kβ localize to vesicle organelles within mouse neurons

Prior studies have demonstrated a role for PI5P4Kα and PI5P4Kβ in clathrin-mediated transport and autophagy (Kamalesh et al., 2017; Lundquist et al., 2018), highlighting a possible vesicular distribution for the PI5P4Ks. To investigate their subcellular localization, we performed double immunofluorescence of PI5P4Kα and PI5P4Kβ with a panel of subcellular markers. By immunofluorescence, we had difficulty detecting endogenous PI5P4Kα and PI5P4Kβ expression. Therefore, we used mCherry-tagged constructs for both PI5P4Kα and PI5P4Kβ. We overexpressed mCherry-tagged PI5P4Kα (Fig. 10A-D) or mCherry-tagged PI5P4Kβ (Fig. 11A-D) in neurons for this analysis

**Figure 10.**
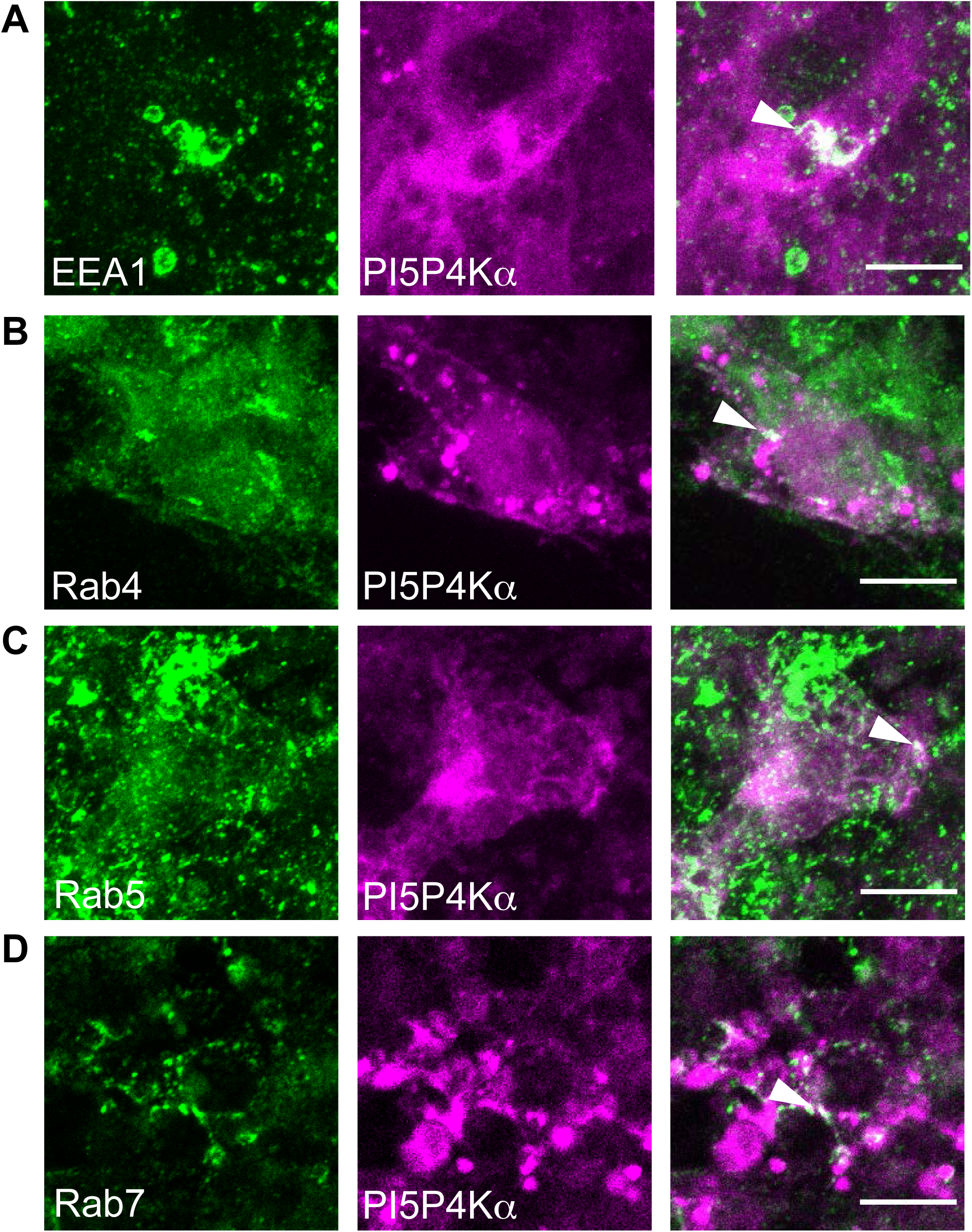
Subcellular distribution of PI5P4Kα in cortical neurons PI5P4Kα co-localizes with the early endosome marker, EEA1 (A), and partially co-localizes with the small GTPases, Rab4 (B), Rab5 (C), and Rab7 (D). Arrowheads point to co-localization. Scale bars = 10 μm.

**Figure 11.**
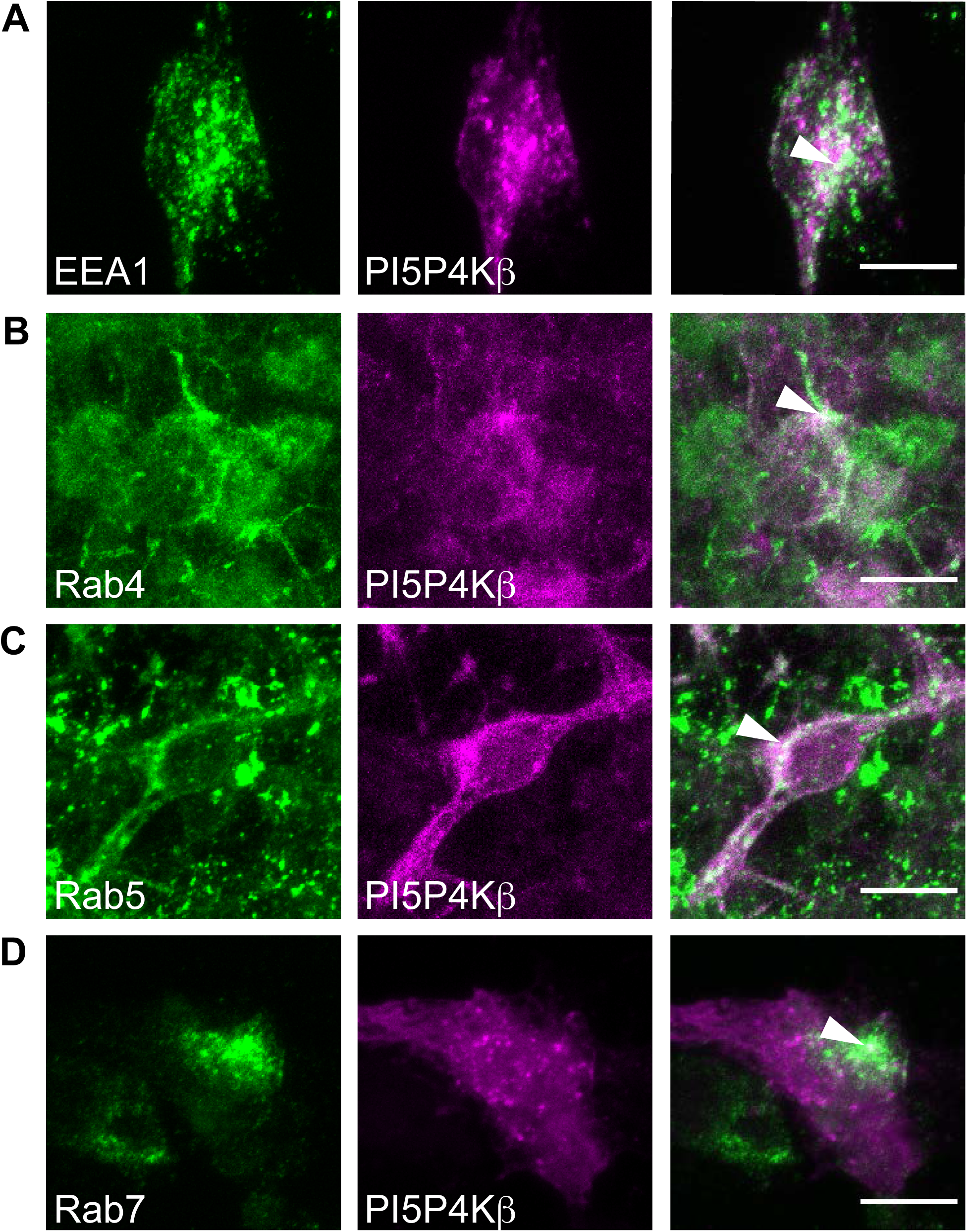
Subcellular distribution of PI5P4Kβ in cortical neurons PI5P4Kβ co-localizes with the early endosome marker, EEA1 (A), and partially co-localizes with the small GTPases, Rab4 (B), Rab5 (C), and Rab7 (D). Arrowheads point to co-localization. Scale bars = 10 μm.

Because PI5P4K*γ* was shown to localize to a vesicular compartment in neurons and HeLa cells (Clarke et al., 2009), we focused on intracellular vesicular pathways for our analysis. Similar to the prior report on PI5P4K*γ*, we found that PI5P4Kα and PI5P4Kβ co-localized with the early endosome marker, EEA1. To further explore the role of PI5P4Kα and PI5P4Kβ in vesicular trafficking, we investigated their co-localization with the small GTPases, Rab4, Rab5, and Rab7, which function in various aspects of intracellular vesicle processing. We found that PI5P4Kα and PI5P4Kβ partially co-localized with Rab4, Rab5, and Rab7, indicating that not only PI5P4K*γ*, but also PI5P4Kα and PI5P4Kβ, may assist in intracellular vesicle trafficking pathways.

### Ultrastructural localization of PI5P4Kα and PI5P4Kβ in the mouse brain to the synapse

Given that our study of subcellular PI5P4Kα and PI5P4Kβ distribution was accomplished through the over-expression of tagged protein, we studied the endogenous localization of PI5P4Kα and PI5P4Kβ using peroxidase and immunogold labeling at the ultrastructural level. We confirmed specificity of our antibodies by quantifying silver-intensified gold (SIG) particles for these proteins in pip4k2a and pip4k2b wild-type and knockout brains. Indeed, we found more SIG particles in PI5P4Kα and PI5P4Kβ wild-type than corresponding knockout brains (Fig. 12A).

**Figure 12.**
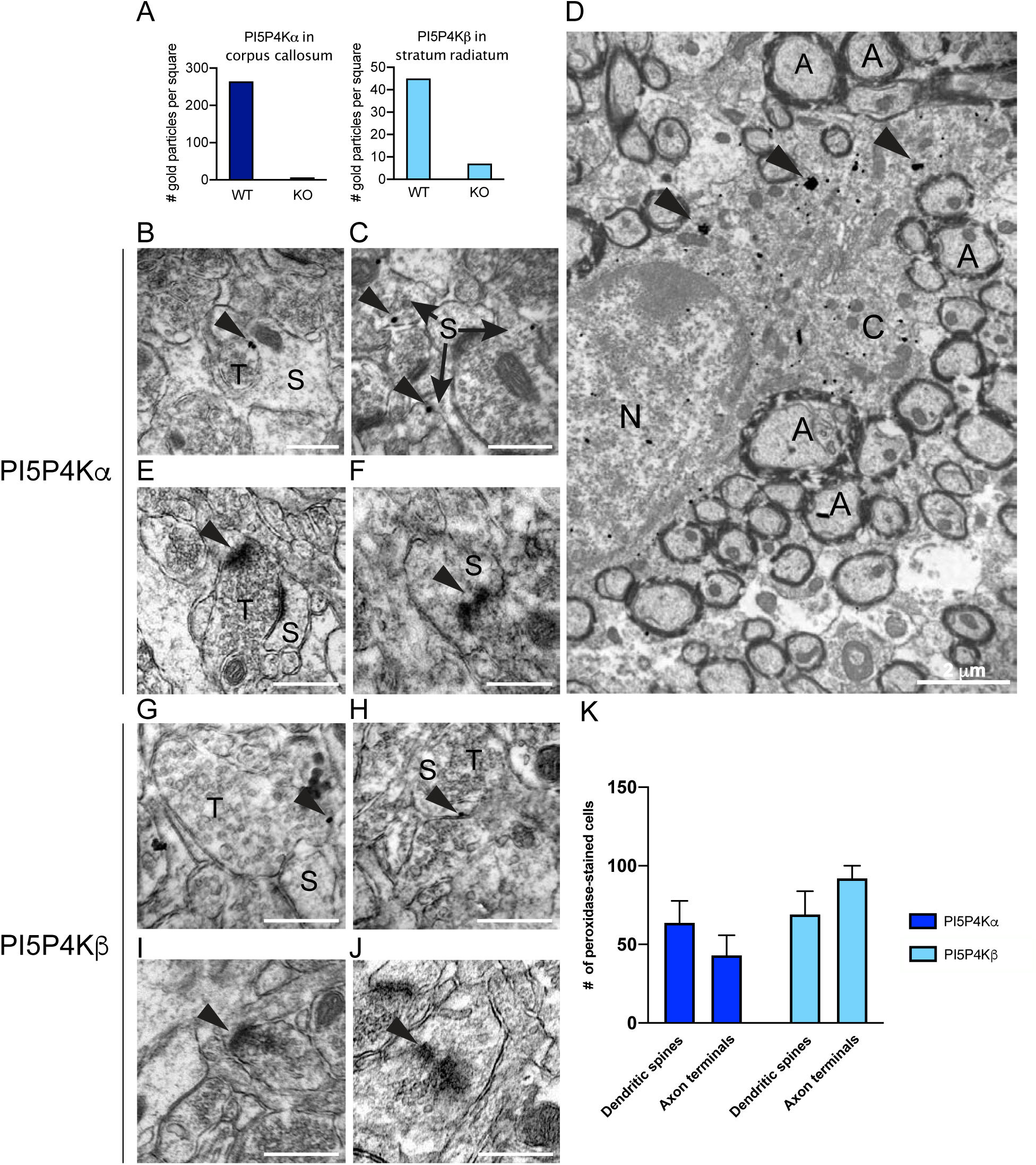
Immuno-electron microscopy of endogenous PI5P4Kα and PI5P4Kβ expression in mouse hippocampus and corpus callosum. A. Quantification of SIG particle labeling for PI5P4Kα in corpus callosum and PI5P4Kβ in stratum radiatum (SR) of hippocampal CA1 from wild-type and respective knockout mice B-D SIG particle labeling for PI5P4Kα is contained in axon terminals (B), dendritic spines in CA1 SR (C), and oligodendrocyte cytoplasm in corpus callosum (D). E-F Immunoperoxidase labeling for PI5P4Kα is contained in axon terminals (E) and dendritic spines (F) in CA1 SR. G-H SIG labeling for PI5P4Kβ is found in axon terminals (G) and dendritic spines (H) in CA1 SR. I-J Immunoperoxidase labeling for PI5P4Kβ in axon terminals (I) and dendritic spines (J). K Quantification of PI5P4Kα and PI5P4Kβ immunoperoxidase labeled profiles in CA1 (n=3 mice per group). ns = not significant Arrowheads point to SIG particles and/or immunoperoxidase labeling. Scale bars for B, C, E, F, G, H, I, J = 500 nm; scale bars for D = 2 μm. A, axon; C, cytoplasm; S, dendritic spine; N, nucleus; T, axon terminal

Because we identified high levels of expression of PI5P4Kβ in the CA1 region of the hippocampus in prior analyses, we decided to characterize the ultrastructural labeling of the PI5P4Ks in this region. PI5P4Kα was localized to axon terminals, with immunogold labeling demonstrating the presence of SIG particles adjacent to the synaptic membrane (Fig. 12B). Similarly, we found PI5P4Kα labeling in dendritic spines, just adjacent to the synaptic membrane (Fig. 12C). Quantification of peroxidase labeled profiles showed that PI5P4Kα was slightly more common in dendritic spines than axon terminals (Fig. 12K). Immunogold analysis also demonstrated ubiquitous PI5P4Kα labeling in the perinuclear region in oligodendrocytes throughout the corpus callosum (Fig. 12D), supporting our light microscopic findings of oligodendrocyte localization.

Similar to PI5P4Kα, PI5P4Kβ localized to both axon terminals and dendritic spines by immunogold (Fig. 12G-H) and peroxidase labeling (Fig. 12I-J). The overall peri-synaptic distribution for both kinases was nearly identical, except that peroxidase quantification demonstrated that PI5P4Kβ tended to be found more often in axon terminals than dendritic spines (Fig. 12K).

## Discussion

In this study, we have characterized the developmental and regional localization of PI5P4Kα and PI5P4Kβ in the mouse brain. Furthermore, we have demonstrated the cell type-specific expression of these kinases in mouse, macaque, and human brain with immunohistochemistry and an independent RNAseq dataset. Finally, we provide the first endogenous expression of PI5P4Kα and PI5P4Kβ by electron microscopy.

### Developmental expression of PI5P4Kα and PI5P4Kβ

Prior work has shown that PIP4K2A and PIP4K2B may play overlapping roles in the stress response after birth. Whereas germline deletion of either pip4k2a or pip4k2b results in normal embryonic growth and development, deletion of both genes results in early embryonic lethality, with pups dying around 12 hours after birth (Emerling et al., 2013). Data from this work showing lethality of tp53^-/-^; pip4k2b^-/-^ embryos indicated that PIP4K2B may be preferentially more important than PIP4K2A shortly after birth. In support of these findings, we demonstrate through immunohistochemistry and RNAseq analysis that PI5P4Kα and PI5P4Kβ display a variable expression pattern in the mouse brain across development. PI5P4Kα is expressed later in development, whereas PI5P4Kβ is expressed at a greater level early in development and diminishes thereafter. It is possible that part of the function of PI5P4Kβ to reduce nutrient stress after birth takes place within the brain and that PI5P4Kβ is less necessary later in development.

### Regional and cell type-specific expression of PI5P4Kα and PI5P4Kβ

Though it is known that the PI5P4Ks are highly expressed in the brain, the distribution of PI5P4Kα and PI5P4Kβ has not been previously described. We find that PI5P4Kα is contained predominantly in white matter with less frequent localization in gray matter and that PI5P4Kβ is expressed solely in gray matter. In the mouse brain, on a cellular level, PI5P4Kα is expressed most notably in oligodendrocytes, though we did observe expression in some neuronal populations. However, PI5P4Kβ is contained solely in neurons. These expression patterns are supported by studies in macaque and human brain tissue.

The restricted localization of the PI5P4Ks to distinct cell types suggests that different cellular processes may regulate their expression. One possibility is that each kinase is expressed through unique transcriptional programs in each cell type. For example, transcription factors, such as the *Olig* family in oligodendrocytes and the *Sox* family in neurons, may regulate expression of each kinase in a cell-autonomous fashion. Alternatively, PI5P4Kα and PI5P4Kβ may be regulated through post-transcriptional or post-translational mechanisms, with trafficking of each kinase to particular intracellular membranes being a highly likely mechanism of localization and potential degradation (Hinchliffe et al., 1999).

The differential localization pattern of PI5P4Kα and PI5P4Kβ highlight potential diverse functions in the brain. For example, PI5P4Kα expression in oligodendrocytes may indicate a potential role in the process of myelination. Interestingly, prior studies have implicated a polymorphism in PIP4K2A (N251S) to inherited risk for schizophrenia (Schwab et al., 2006; He et al., 2007; Rethelyi et al., 2010; Thiselton et al., 2010), with lymphocytes from schizophrenia patients expressing elevated levels of PIP4K2A (Saggers-Gray et al., 2011). Several studies have postulated white matter dysfunction in schizophrenia, arguing for a role for oligodendrocyte dysfunction in this disease (Tonnesen et al., 2018). Therefore, our findings of oligodendrocyte-specific PIP4K2A expression may connect the epidemiological studies with the white matter theory in schizophrenia.

On a molecular level, PI5P4Kα influences several neurotransmitter systems. PI5P4Kα has been shown to increase the expression of the neuronal glutamate transporter, EAAT3, while the schizophrenia-associated polymorphism, N251S, exerts a dominant-negative effect on EAAT3 expression (Fedorenko et al., 2009), resulting in reduced glutamate uptake. PI5P4Kα, but not the N251S mutant, activates neuronal M channels, which under normal physiological conditions, suppress dopaminergic transmission (Fedorenko et al., 2008). Interestingly, this mutant does not alter PI5P4Kα intrinsic kinase activity, suggesting a non-catalytic function of this polymorphism in disrupting PI5P4Kα function (Clarke and Irvine, 2013). Our data showing a peri-synaptic distribution of PI5P4Kα support these studies that demonstrate a possible role of PI5P4Kα in the molecular pathogenesis of schizophrenia.

It is also noteworthy that several genetic susceptibility loci in PI5P4K*γ* have been linked to autoimmune diseases, including rheumatoid arthritis and type 1 diabetes mellitus, and that PI5P4K*γ*-deficient mice exhibit a hyperimmune phenotype (Barton et al., 2008; Fung et al., 2009; Shim et al., 2016). Though there have not been as of yet any PIP4K loci linked to the development of multiple sclerosis, the predilection of PIP4K2A expression to oligodendrocytes may be relevant to demyelinating diseases. Given that both PI5P4Kα and PI5P4Kβ, but not PI5P4K*γ*, exhibit kinase activity, it is possible that their differential expression in oligodendrocytes and neurons, respectively, highlights non-overlapping kinase function in these cell types. In addition, the similar pattern of expression of PI5P4Kβ and PI5P4K*γ* may indicate that either PI5P4K*γ* is required for appropriate localization of PI5P4Kβ or that these 2 kinases serve unique functions in neurons. For example, germline deletion of pip4k2c in mice results in a dramatic increase in S6-Kinase activity downstream of TORC1 in the mouse brain (Shim et al., 2016). Such upregulation of S6-Kinase activity may be explained by the role of PI5P4K*γ* in inhibiting PI4P5K activity and downstream PI 3-kinase activity (Wang et al., 2019). Therefore, PI5P4Kα and PI5P4Kβ kinase activity could balance PI4P5K*γ* inhibitory activity in a cell-autonomous manner.

### Subcellular distribution of PI5P4Kα and PI5P4Kβ

The localization of PI5P4Kα and PI5P4Kβ to the early endosome system and to the perisynaptic region continues to support the role of the PI5P4Ks in intracellular PI-4,5-P_2_ pathways (Clarke et al., 2009). At a subcellular level, each kinase exhibits similar co-localization with markers of the early endosome system, and likewise, ultrastructural analysis demonstrates that PI5P4Kα and PI5P4Kβ are both found in peri-synaptic regions in axon terminals and dendritic spines. While it is possible that these kinases participate in ubiquitous vesicle trafficking, their particular localization adjacent to the synaptic membrane in both axon terminals and dendritic spines suggests that they may hold a more specific function in the brain, namely synaptic neurotransmission. Given that we did not identify their specific localization on the synaptic membrane, it is perhaps more likely that these kinases serve a role in the synaptic vesicle docking or recycling process rather than vesicle fusion or release at the synapse.

### Conclusions

To our knowledge, this is the first description of the endogenous ultrastructural localization of the PI5P4Ks in the brain. Their differential expression pattern in the brain highlights potentially diverse functions of these kinases, and their novel localization to the peri-synaptic region within particular axon terminals and dendritic spines lends support to the theory that these kinases may serve a supplementary role in synaptic function, whether through synaptic vesicle docking or recycling processes.

## Acknowledgements

Neuroanatomy EM Core; Dr. Jonathan Victor and Dr. Keith Purpura (supported by NIH EY9314) for the donation of the macaque tissue.

L.C.C. is a founder and member of the BOD and SAB of Agios Pharmaceuticals; he is also a co-founder, member of the SAB, and shareholder of Petra Pharmaceuticals. These companies are developing novel therapies for cancer. L.C.C. laboratory receives some funding support from Petra Pharmaceuticals.

The data that support the findings of this study are available from the corresponding author upon reasonable request.

